# Fucoidan from brown algae *Laminaria japonica* trigger apoptosis in colon cancer cells

**DOI:** 10.64898/2026.05.31.728131

**Authors:** Koya Shimabukuro, Ikuko Miyagi, Nanae Harashima

## Abstract

In many cases, cancer cells develop resistance to chemotherapy and other cancer treatments. Therefore, there is a need for new therapeutic agents using naturally derived compounds that are expected to have low toxicity and fewer side effects. Fucoidan is a sulfated polysaccharide found in brown algae such as kelp and wakame seaweed. Many previous reports have shown that fucoidan exerts anti-bacterial, anti-viral, antioxidant, immunomodulatory effects, and anti-tumor effects. The antitumor and antiviral effects of fucoidan have been reported to vary depending on its origin, as they are influenced by sulfate content and molecular weight. Therefore, it is important to investigate the antitumor effects of various species of fucoidan, but there are few reports on the effects of fucoidan derived from *Laminaria japonica* on colorectal cancer. In this study, we evaluated the effects of fucoidan from *Laminaria japonica* on apoptosis in five human colon cancer cells. The apoptotic cell population was significantly increased in fucoidan-treated cells. In addition, the expressions of Bax, Bak, PARP, caspase-8, -9 and -3 were upregulated. The necroptosis-related molecule RIP and MLKL were degraded indicates that necroptosis was not involved in this fucoidan-treated cell death. These results suggest that fucoidan-treated cells showed induction of apoptosis via mitochondrial intrinsic pathway, but not necroptosis via caspase-8. Fucoidan-induced apoptosis may prove useful in the therapeutic protocol of colon cancer.

## Introduction

Colorectal cancer is the third most common cancer worldwide, and the number of new cases globally is projected to reach 3.2 million by 2040 (1). Advances in the pathophysiology of colorectal cancer have led to an expansion of treatment options, including existing chemotherapy, surgery, radiation therapy, and immunotherapy. Early-stage colorectal cancer has high treatment efficacy and survival rates. However, the effectiveness of these treatments is limited for advanced cancer, creating a need for personalized medicine and a better understanding of the mechanisms underlying treatment resistance.

Research on bioactive compounds derived from marine sources and seaweed extends not only nutritional aspects but also investigation of medical and therapeutic applications. In cancer treatment research, several bioactive compounds from marine sources have shown potential as apoptosis-inducing agents, and clinical trials are currently underway. Bioactive compounds identified in brown algae include fucoxanthin, fucoidan, laminarin, and sargaquinoic acid (2). Fucoxanthin is a carotenoid, a naturally occurring pigment that contains both polar hydrocarbon carotene and polar compounds known as xanthophylls (3). Both fucoidan (4, 5) and laminarin (6) are polysaccharides derived from brown algae, but fucoidan is a type of sulfated polysaccharide and is widely used in both industrial and medical applications. Various molecular mechanisms have been reported for these compounds, including the induction of apoptosis, anti-migratory effects, and DNA damage in various cancer cells.

Fucoidan is one of essential anticancer macromolecules with low toxicity and few side effects. Due to its wide range of physiological activities, including immune modulation, antitumor, antiviral, and blood glucose-lowering effects, it has recently garnered attention as a potential therapeutic candidate (7). The biological activity of fucoidan is believed to be influenced by its sulfate content and molecular weight, which depends on the species, harvest time, and extraction method (8). Therefore, it is important to investigate the anti-tumor effects of fucoidan from various species. Previous reports have described the anti-tumor effects of fucoidan derived from bladderwrack (*Fucus vesiculosus*) on human colorectal cancer cells via Akt signaling (9) and apoptosis induction (10), as well as the ROS-dependent JNK activation-mediated apoptosis induction of fucoidan derived from mozuku (*Cladosiphon novae*-*caledoniae Kylin*) from the Kingdom of Tonga on breast cancer cells (11). However, there are limited reports on the effects of fucoidan derived from *Laminaria japonica* on human colorectal cancer cells. The biological activity of fucoidan is believed to be influenced by its sulfate content and molecular weight, which depends on the marine species, harvest time, and extraction method (8). Therefore, it is important to investigate the antitumor effects of fucoidan from various species. In this study, we investigated the cell death-related molecule expression and anti-cancer effects of fucoidan from *Laminaria japonica* on human colorectal cancer cells. Our data indicate that fucoidan inhibited cancer cell growth, migration, and promoted apoptotic cell death.

## RESULTS

### Fucoidan inhibited the proliferation of colorectal cancer cells

To investigate the effects of fucoidan on colorectal cancer cell lines, cell viability was assessed using the WST-8 assay. In cultures treated with 800–1200 µg/mL fucoidan, a significant concentration- and time-dependent decrease was observed in all five tested cell lines (Figure 1). For subsequent experiments, the final concentration of fucoidan was set as follows; for SW480 and SW620 were 1000 μg/mL, for DLD-1, LS174T and HT-29 were 800 μg/mL.

**Figure 1.**
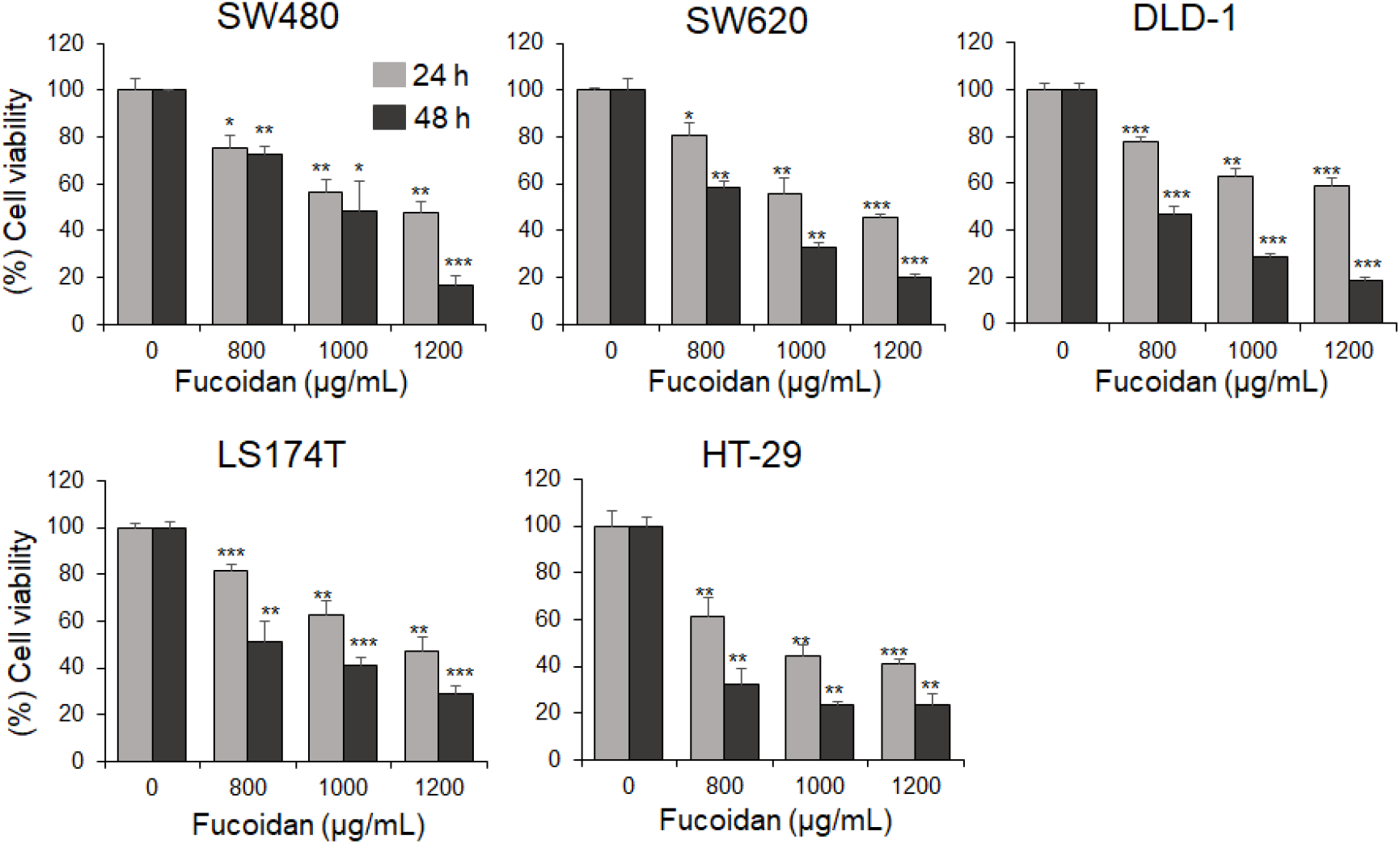
Effects of fucoidan on cell viability of colorectal cancer cells. Cells were treated with the indicated concentrations of fucoidan for 24 h or 48 h. The cell viability was evaluated by WST-8 assay. Data are expressed as the mean standard deviation (means ± SD) of three independent experiments with samples in triplicate. \**p* < 0.05, \*\**p* < 0.01 versus control.

In cells treated with fucoidan, the gaps between cells widened, cell adhesion decreased, and cells began to detach, with blob-like cells and fragmented cells observed (Figure 2A). Cell adhesion decreased 1 hour after addition, blebbing cells were observed 12 hours after addition, and fragmented cells were observed more than 24 hours after addition treatment (Figure 2A). Colonies form when a single cell divides indefinitely (Figure 2B). Assessing colony formation capacity is crucial for investigating the uncontrolled division characteristics of cancer cells.

**Figure 2.**
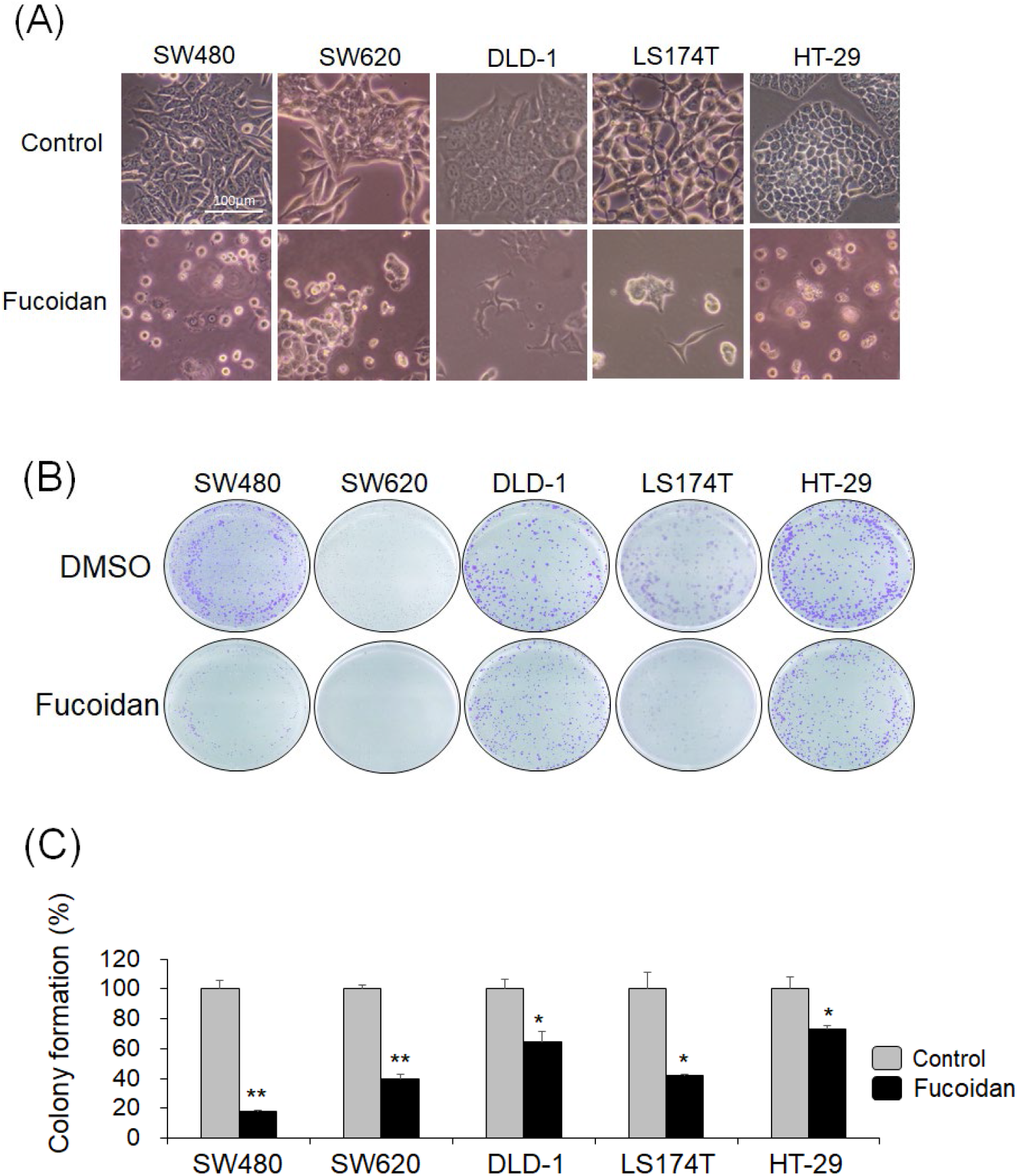
Effects of fucoidan on cell proliferation of colorectal cancer cells. (A) Cells were treated with fucoidan for 48 h. Cell density and morphological changes were visualized using a phase contrast microscope (magnification, x 200). (B) Colony images stained with crystal violet. (C)The number of colonies per well were counted and calculated. The percentages of colonies are expressed as the mean standard deviation (means ± SD) of three independent experiments with samples in triplicate. \**p* < 0.05, \*\**p* < 0.01 versus control.

When the number of colonies in the solvent control cells was set to 100%, the results showed a significant reduction in the number of cell colonies following fucoidan treatment (Figure 2C). Particularly colony formation was inhibited by more than 60% in the SW480 and SW620 cell lines, indicating that fucoidan exerts a strong antiproliferative effect on these cancer cells.

### Fucoidan inhibited the migration of colorectal cancer cells

Wound healing assay was performed to examine the invasive capacity of cells treated with fucoidan. The invasive capacity of cancer cells is crucial for evaluating metastasis. The black areas in the image represent the regions scratched by the tip, while the adjacent white areas indicate the cells (Figure 3A). In cells treated with fucoidan, the invasion rate was significantly lower compared to cells in the solvent control. The migration closure rates for each colorectal cancer cell line were 23.6 ± 7.3% for SW480 cells, 2.9 ± 3.7% for SW620 cells, 24.5 ± 3.3% for DLD-1 cells, and 3.2 ± 3.6% for HT-29 cells (Figure 3B). The migration closure rates in the fucoidan-treated groups were significantly lower than those in the control groups. These results clearly indicate that fucoidan can inhibit the ability of colorectal cancer cells to migrate and invade and therefore metastasize the secondary locations.

**Figure 3.**
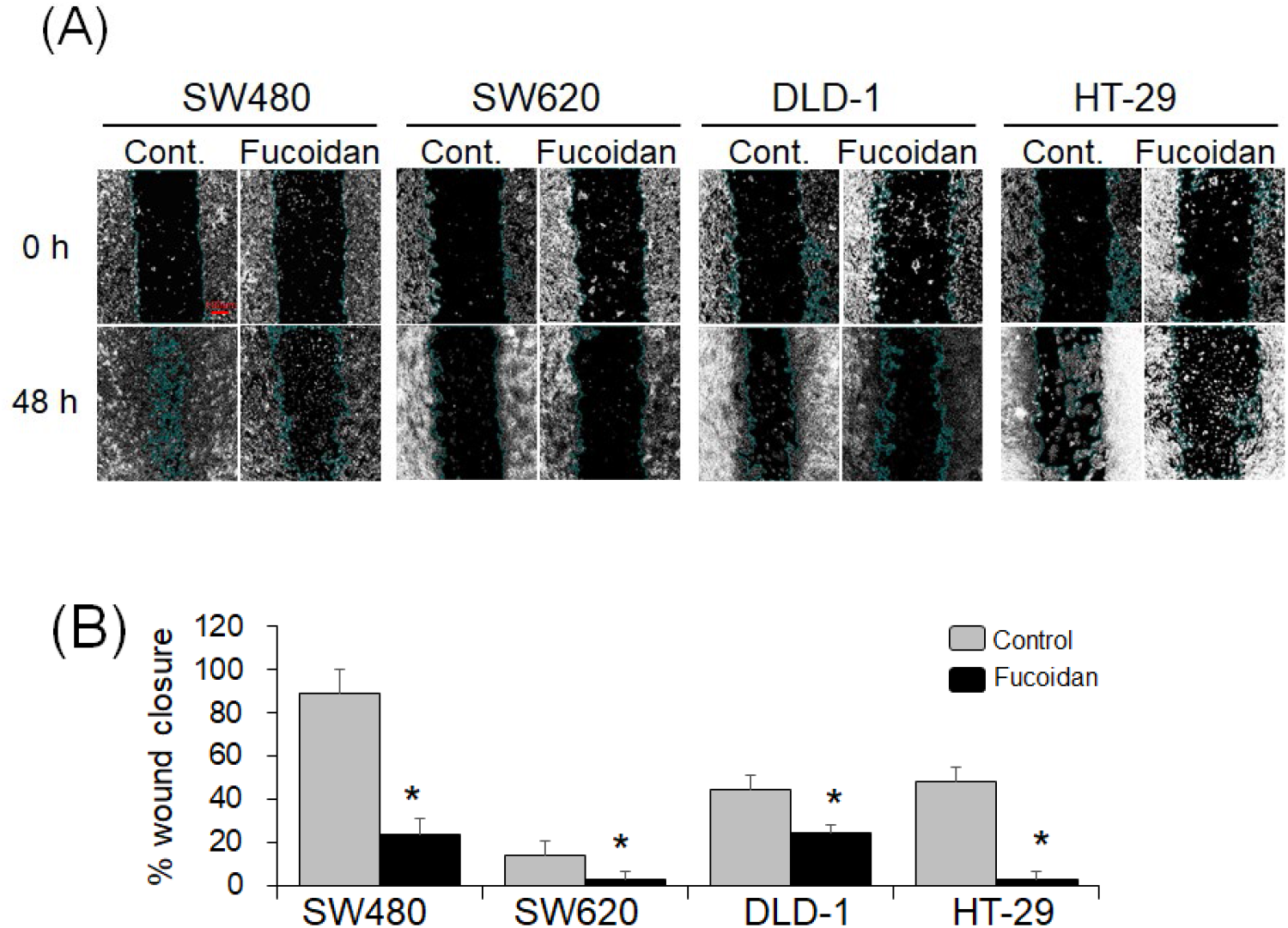
Effects of fucoidan on colorectal cancer cell migration by wound healing assay. (A) Cells were seeded into culture plates and grown to 90-100% confluence. The cells were scratched with a sterile tip, fucoidan (0, 800 or 1000 μg/mL) were added and wounded areas were photographed at 0, 24, and 48 h. (B) The inhibitory effect on cell migration is expressed as the mean percentage of wound closure in triplicate. A significant difference from the vehicle was indicated as \**p* < 0.05.

### Fucoidan induces apoptosis in colorectal cancer cells

AO penetrates the cell membranes of both live and dead cells, staining all cells and producing green fluorescence. In contrast, PI penetrates only cells with damaged cell membranes, staining dead cells and producing red fluorescence. The degree of staining with PI varies depending on the stage of cell death, with dead cells staining yellow, orange, or red (Figure 4A). In the cells cultured for 48 hours after fucoidan treatment, the number of dead cells significantly increased compared to the control cells (Figure 4B).

**Figure 4.**
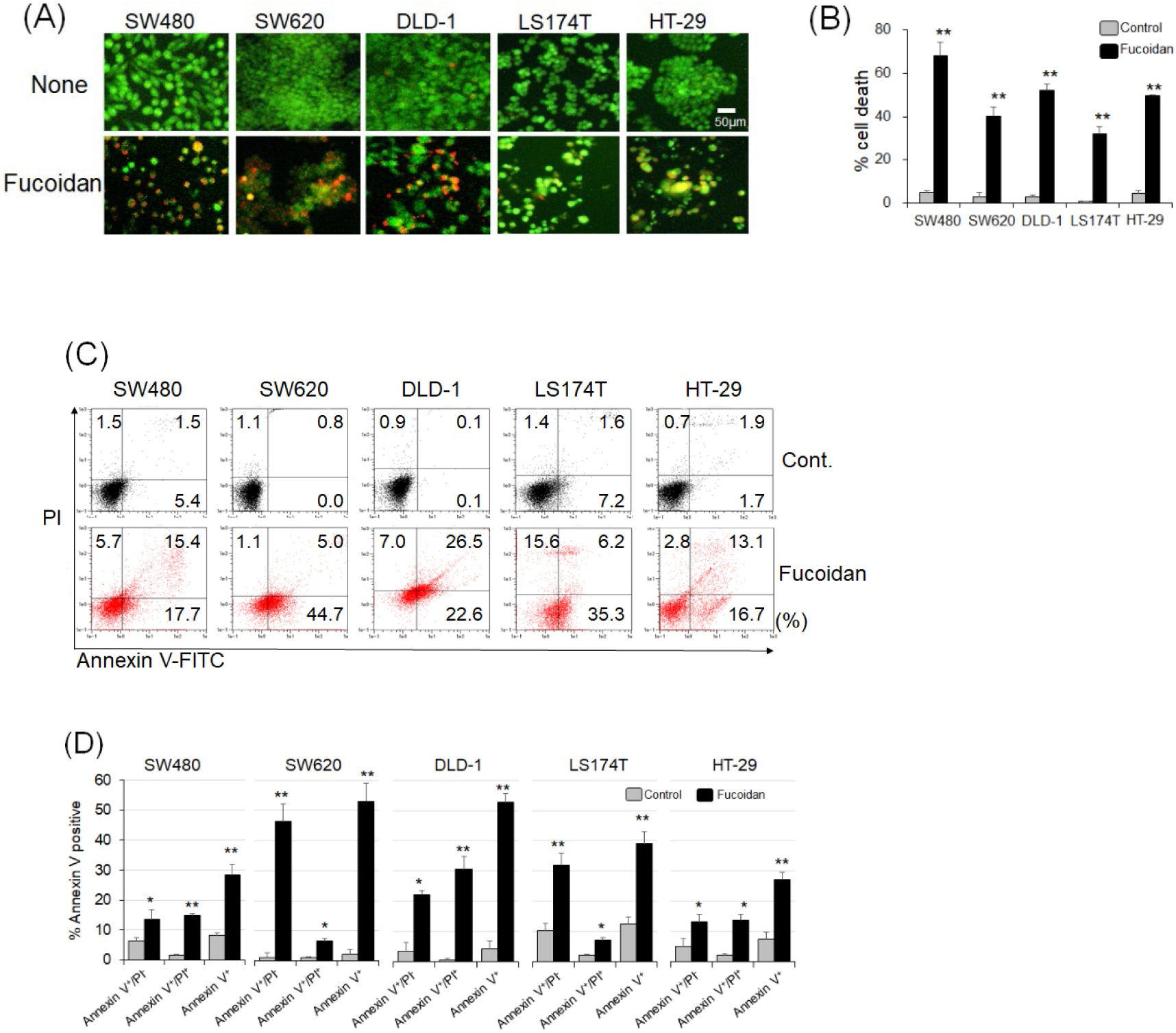
Fucoidan induces apoptosis in colorectal cancer cells. (A) Cells were treated with fucoidan for 48 h. Acridine orange/propidium iodide double staining to evaluate for cell death (magnification, x 200). Images show the morphological features and stages of viable (green), earl phase of apoptosis (yellow to orange) and late phase of apoptosis/necrotic (orange to red) in cells treated with fucoidan. Scale bar represents 50 μm. (B) Bar diagrams show percentage of apoptotic cells (yellow to red). The percentages of cell death are expressed as the means ± SD of three independent experiments, each performed in triplicate. (C)Various stages of cellular apoptosis were detected by flow cytometry. (D) Bar diagrams show percentage of apoptotic cells (Annexin V positive and/or PI positive). Percentages of cell death are expressed as the means ± SD of three independent experiments, each performed in triplicate. \*\**p* < 0.01 versus control.

Apoptosis analysis was performed using flow cytometry to assess cell death trends. In the dot plot generated by double staining with Annexin V-FITC and PI, the lower-right and upper-right quadrants represent early and late apoptosis, respectively (Figure 4C). The proportion of apoptotic cells significantly increased in all colorectal cancer cells treated with fucoidan (Figure 4D). Early and late apoptotic cells were observed in SW480, DLD-1, and HT-29 cells, however, fucoidan treatment primarily induced early apoptosis in SW620 and LS174 cells. It was found that the induction of apoptosis varied depending on the cell line used.

Mitochondria are one of the key organelles in the body when discussing early-stage cytotoxicity, oxidative stress, and apoptosis. Mitochondria produce the energy required for biological processes by utilizing oxygen to synthesize ATP, and a decline in mitochondrial activity or dysfunction is closely associated with conditions such as cancer and aging. JC-1 staining was performed to investigate changes in mitochondrial membrane potential (ΔΨm) levels in colorectal cancer cells treated with fucoidan. When ΔΨm is high, the JC-1 dye aggregates within the mitochondria and emits red fluorescence, however, when mitochondrial depolarization occurs and ΔΨm decreases, red fluorescence decreases and green fluorescence is emitted. Ratio of red/green fluorescence decreased in tested cancer cells after treatment with fucoidan, indicating a decrease in ΔΨm (Figure 5B). Particularly ΔΨm was reduced by more than 40% in the SW480, DLD-1 and HT-29 cell lines, indicating that fucoidan promotes apoptotic cell death through mitochondrial pathway in colorectal cancer cells.

**Figure 5.**
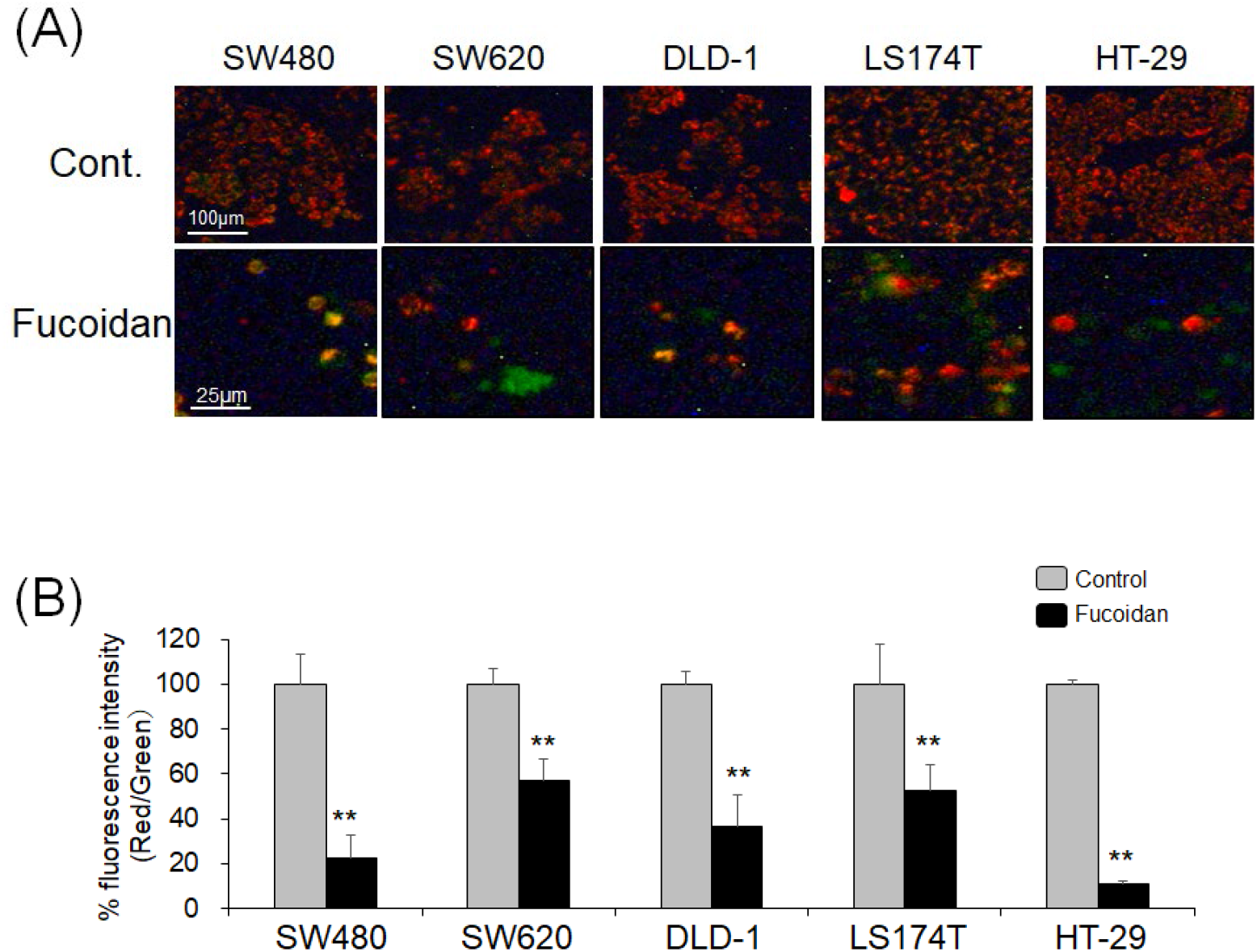
Fucoidan triggers the depolarization of ΔΨm in colorectal cancer cells. (A) JC-1 staining detected changes in mitochondrial membrane potential through fucoidan treatment (magnification, x 200). (B) The red to green fluorescence intensity of JC-1 ratio was significantly decreased in fucoidan-treated cancer cells compared with the control (\*\**p* < 0.01 versus control). The percentages are expressed as means ± SD of three independent experiments with samples in triplicate.

### Effect of fucoidan on the expression of extrinsic- and intrinsic-apoptosis

There are two pathways of apoptotic cell death: the intrinsic pathway and the extrinsic pathway. We examined the protein expression of apoptosis- and necroptosis-related molecules in colorectal cancer cells co-cultured with fucoidan (Figure 6). The Bid expressions were reduced in all cancer cell lines, and the expression of tBid, resulting from Bid activation, was observed in SW480, SW620, and DLD-1(Figure 6A). The activation of caspase-8 detected as cleaved form was showed in SW480, SW620, DLD-1, and HT-29 cells (Figure 6A). These results indicated that fucoidan induced apoptosis via extrinsic pathway including caspase-8 activation. Fucoidan increased Bax expression in SW480 and HT-29 cells, increased Bak in LS174T and HT-29 cells, and decreased Bcl-2 expression in SW620 and LS174T cells (Figure 6B). Activated caspase-9 (cleavage forms) and PARP were observed in all tested cell lines, activated caspase-3 were observed in SW480, SW620, and DLD-1 (Figure 6B).

**Figure 6.**
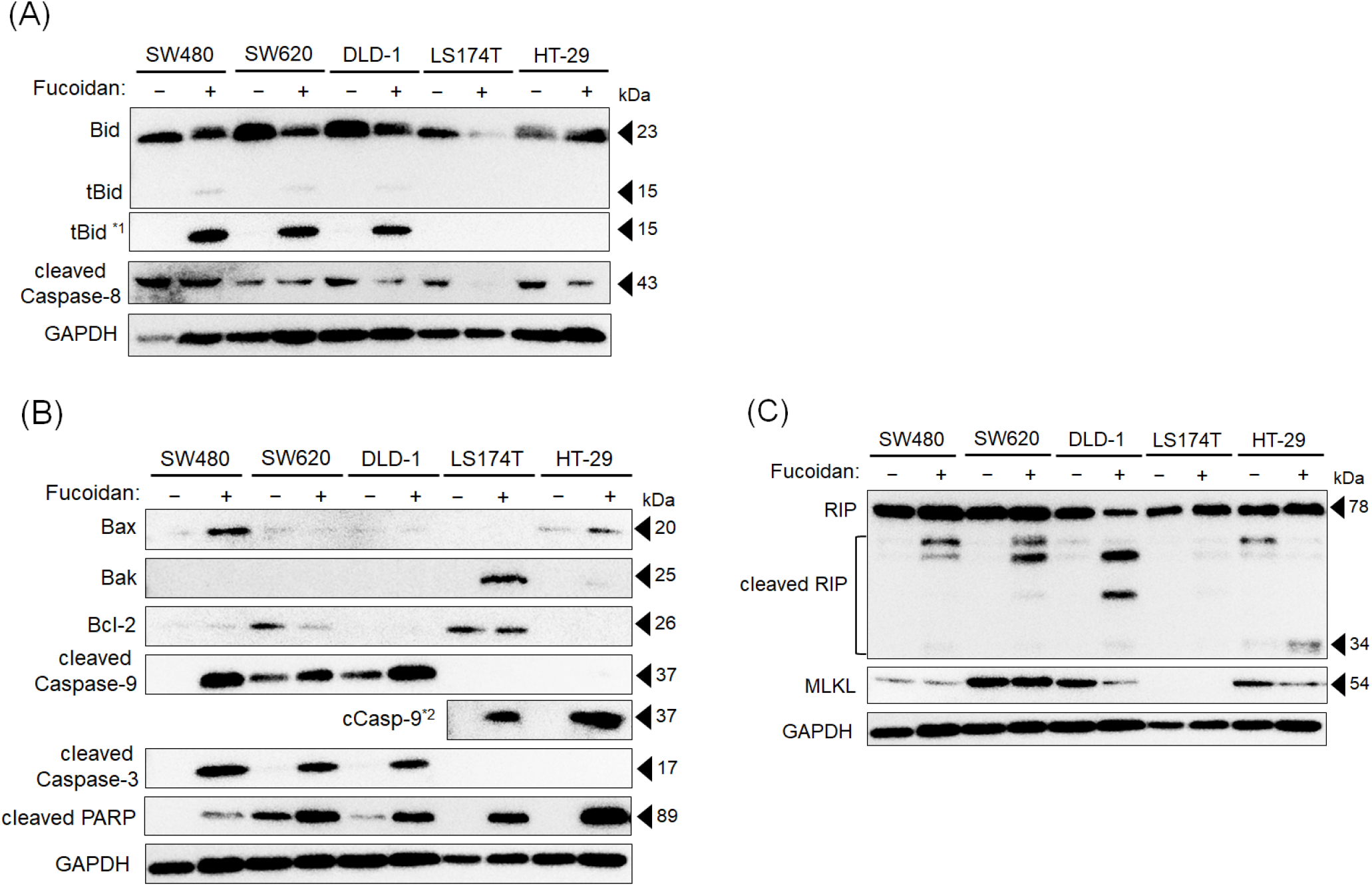
Fucoidan modulates the expression levels of apoptosis-related molecules in colorectal cancer cells. Cells were treated with fucoidan for 48 h. The protein levels of indicated genes were detected by western blotting analysis. The expressions of Bid and caspase-8 (A), Bax, Bak, Bcl-2, caspase-9/-3, and PARP cleavage forms (B), and RIP and MLKL (C) were detected in fucoidan-treated colorectal cancer cells. Exposure time was extended to ^*1^5 min or ^*2^4 min.

During apoptosis, caspase-8 suppresses necroptosis by cleaving and inactivating RIP (12-14). MLKL is a necroptosis executor activated by the RIP-RIP3 complex (15). Cleavage of RIP was detected in SW480, SW620, DLD-1, and HT-29 cells, which exhibited caspase-8 activation (Figure 6C). Furthermore, the expression of MLKL decreased in SW620, DLD-1, and HT-29 cells (Figure 6C). These results suggest that fucoidan suppressed necroptosis and modulated both extrinsic- and intrinsic-pathway of apoptosis in colorectal cancer cells.

## DISCUSSION

Fucoidan reduced the cell viability of human colorectal cancer cells in a concentration- and time-dependent manner (Figure 1). Although the species differ, fucoidan derived from the *Sargassum genus* did not reduce cell viability in 293T cells (human fetal kidney epithelial cells) at a final concentration of 800 µg/mL after 48 hours of incubation, therefore it is considered that fucoidan does not affect normal cells (8). Fucoidan treated ccolorectal cancer cells were observed to detach and float immediately after addition (Figure 2A). This is thought to be due to fucoidan inhibiting adhesion molecules such as integrins present on the surface of tumor cells (16). Inhibition of integrin binding triggers the cleavage of caspase-8, leading to the induction of apoptosis (17).

There are two pathways of apoptotic cell death, the extrinsic and intrinsic pathway. The extrinsic pathway is initiated by the interaction between cell surface death receptors which belong to the tumor necrosis factor receptor (TNFR) superfamily, and their respective TNF family ligands. Signal transduction involving death receptors and their ligands includes TNFR1-TNFα, FAS-FASL, and DR4/DR5-TRAIL. Upon ligand binding to the death receptor, caspase-8 binds via an intracellular adaptor molecule called Fas-associated via death domain (FADD), forming a complex (DISC). As a result, caspase-8 is activated, leading to the activation of caspase-3, an apoptotic executor (18). Bid, a BH-3-only protein of the Bcl-2 family, is cleaved into tBid by activating caspase-8, which then activates Bax and Bak (19). The intrinsic pathway is a mitochondria-mediated apoptotic cell death pathway. Non-receptor-mediated stimuli, such as chemical stimuli, radiation, and stress, converge on the mitochondria to generate intracellular signals (16). One such mechanism involves regulation by p53. When DNA damage occurs due to stimulation, activation of the tumor suppressor protein p53 induces the transcription of pro-apoptotic effector molecules of the Bcl-2 family, such as Bax and Bak, while suppressing the transcription of anti-apoptotic molecules such as Bcl-2 (17). The activation of Bax and Bak forms pores in the mitochondrial outer membrane, increasing mitochondrial outer membrane permeability (20). When mitochondrial membrane permeability increases, cytochrome c is released from the mitochondria, forming an apoptosome in association with Apaf-1 and caspase-9, which stimulates the activation of caspase-9. Activated caspase-9 activates the apoptotic effector caspases-3/6/7, thereby inducing apoptosis (21). The DNA repair molecule PARP is cleaved by caspase-3, and PARP cleavage is a hallmark of apoptosis (22).

Since pro-apoptotic molecules and caspases were upregulated and anti-apoptotic molecules were downregulated, it is considered that fucoidan induced both intrinsic and extrinsic apoptosis in colorectal cancer cells (Figure 6). In addition, necroptosis-related molecules, such as RIP and MLKL, were degraded in fucoidan-treated cancer cells (Figure 6C). These results suggest that fucoidan more strongly induces apoptosis and ultimately causes cell death by suppressing or degrading necroptosis-related molecules. LS174T and HT-29 cells did not show activation of caspase-3, an apoptosis execution factor. However, PARP degradation was observed, suggesting that apoptosis was occurring, finally. There are reports that caspase-7 activation occurs more readily than caspase-3 activation in HT-29 cells at low concentrations of additives (23). Therefore, it is thought that PARP degradation and apoptotic cell death are carried out not by caspase-3, but by other apoptosis execution factors such as caspase-7.

According to ATCC cell line information, SW480 and LS174T represent stage II colorectal cancer, while SW620 and DLD-1 represent stage III colorectal cancer. Colorectal cancer cells with lower cancer stages (SW480, LS174T) tended to show a higher rate of migration inhibition by fucoidan (Figure 3). Furthermore, when comparing primary SW480 and metastatic SW620 cells from the same patient, SW480 cells tended to show higher sensitivity to fucoidan. Fucoidan sensitivity varies among cells, but it is thought to be effective against all tested colorectal cancer cells. Cancer staging is intended to predict patient prognosis and select treatment options, and among these, the 5-year survival rate decreases as the stage increases. This is because as the stage increases, cancer metastasizes lymph nodes and other organs, spreading more widely throughout the body and making treatment more difficult. We believe that the possibility of treatment resistance at the cellular level, as revealed in this study, may bring a new perspective to cancer staging and prognosis prediction.

## CONCLUSIONS

Fucoidan derived from *Laminaria japonica* reduced cell viability in dose-dependent manner in five colorectal cancer cell lines examined in this study, suggesting that it induces both extrinsic- and intrinsic-apoptosis. Furthermore, although the sensitivity of *Laminaria japonica*-derived fucoidan differed among cell lines, even among colorectal cancer cells, fucoidan showed a significant anti-cancer effect against examined cell lines, suggesting that fucoidan-sensitivity may vary depending on the stage of colorectal cancer. Further investigations, including *in vivo* experiments, are needed to examine its effects on cancer regression, metastasis suppression, and side effects.

## MATERIALS AND METHODS

### Cell lines and reagents

Five human colorectal cancer cell lines (SW480, SW620, DLD-1, LS174T, and HT-29) were kindly provided by Dr. Keizo Takenaga (Chiba cancer center). All cells were cultured in D-MEM supplemented with 10% fetal bovine serum and 1% penicillin-streptomycin at 37°C and 5% CO_2_. Fucoidan derived from *Laminaria japonica* (Biosynth, UK) was dissolved in dimethyl sulfoxide (DMSO), and a 30 mg/mL stock solution was used for each experiment.

### Cell viability and cell morphology

Human colorectal cancer cells were seeded at 5 × 10^3^ cells/well in 96-well plates. After overnight culture, Fucoidan (concentration: 800 to 1200 μg/mL) was added. After an additional 24 or 48 hours of culture, cell viability was measured at 450 nm using the Cell Counting Kit-8 (DOJINDO, Japan), and cell viability was calculated using cells treated with the solvent control as a reference. Analysis of cell morphological changes by a phase contrast microscope. 5 × 10^4^ cells were seeded in 24-well plates and treated with fucoidan (1000 μg/mL for SW480 and SW620 cells, 800 μg/mL for DLD-1, LS174T and HT-29 dells) for 48 hours. The plates were observed under a phase contrast microscope (CKX53, Olympus).

### Colony formation assay

Human colorectal cancer cells were seeded at 500 cells/well in 6-well plates and cultured overnight. After adding fucoidan and culturing for 48 hours, the medium was replaced with fresh medium, and the cells were cultured for an additional 8 to 12 days. Following fixation, the cells were stained with 0.5% crystal violet solution, and the number of colonies were counted using ImageJ software.

### Acridine orange/propidium iodide (AO/PI) double staining

Human colorectal cancer cells were seeded at 5 × 10^4^ cells/well in a 24-well plate and cultured overnight. Fucoidan was then added, and the cells were cultured for 48 hours. AO (Cellstain AO, DOJINDO)/PI (Cellstain PI, DOJINDO) double staining was performed, and the samples were observed under a fluorescence microscope. One hundred cells per well were counted, and the average value across three wells was calculated to determine the percentage of dead cells.

### Scratch wound healing assay

Colorectal cancer cells were seeded at 4 × 10^6^ cells/well in 24 well culture plates and allowed to attach overnight. Once cells reached 90-100 % confluency, the cell monolayer was scratched with a 200μL pipette tip held vertically followed by washing with PBS to remove floating cells. The cells were treated with fucoidan or complete DMEM medium with DMSO as a control. The wound area was imaged on 0- and 48-hours incubation using a phase-contrast microscope (CKX53, Olympus) connected to a digital camera (Visualix STD2, Visualix). The wound area of the scratched region was calculated using ImageJ (NIH, United States). %Wound closure=A_0h_−A_48h_/A_0h_×100

### Mitochondrial membrane potential (ΔΨm)

Mitochondrial membrane potential was assessed by JC-1 reagent (MitoMP Detection Kit, DOJINDO), following manufacturer’s protocol. Briefly, cancer cells were cultured in 96-well plate (5 × 10^3^ cells/well) and treated with fucoidan for 48 hours. The cells were labelled with JC-1 dye for 60 min and washed with PBS. Fluorescent intensity was measured (ex: 485 nm and em: 535 nm) using a microplate reader (SpectraMax^™^ iD5 and SoftMax Pro, Molecular Devices) and visualized using fluorescent microscope.

### Annexin V/PI analysis by flow cytometry

Human colorectal cancer cells were seeded at 2 × 10^6^ cells/well in 6-well plates and cultured overnight. Fucoidan was added, and the cells were cultured for 48 hours. The cells were then stained with Annexin V-FITC (Nacalai Tesque, Japan) and PI and analyzed using a flow cytometer (MACSQuaify™) and MACSQuaify™ software.

### Western Blotting

Five types of human colorectal cancer cells were seeded at 2 × 10^6^ cells/well in 100 mm dishes and cultured overnight. Fucoidan was added, and after 24 or 48 hours of culture, the cells were harvested and proteins were extracted using M-PER Reagent (PIERCE). Proteins were separated by SDS-PAGE electrophoresis using 10–12.5% acrylamide gels, and Western blotting was performed. The following primary antibodies were used: Bax (Cell Signaling Technology; CST, 5023), Bak (CST, 12105), Bcl-2 (Bio-Legend, 658702), Bid (CST, 2002), cleaved Caspase-9 (CST, 9915), Caspase-8 (GeneTex, GTX110723), cleaved Caspase-3 (CST, 9668), cleaved PARP (CST, 5625), RIP (CST, 3493), MLKL (CST, 14993), and GAPDH (abm, G043) and as secondary antibodies, we used Peroxidase-conjugated AffiniPure Rabbit Anti-Mouse IgG (Jackson ImmunoResearch, 124878) and Peroxidase-conjugated AffiniPure Goat Anti-Rabbit IgG (Jackson ImmunoResearch, 93843).

## Statistical Analysis

Student’s t-test was used to compare groups, with \**p* < 0.05, \*\**p* < 0.01, and \*\*\**p* < 0.001 indicating statistical significance. All experiments were repeated at least 3 times with triplicates treatment. Representative data is shown.

## Authors’ contribution

K. S.: Conceptualization, Methodology, Data curation, Investigation, and Writing. I. M.: Validation, Methodology. N. H.: Conceptualization, Supervision, Resource, Writing-review & editing, Funding acquisition.

## REFERENCES

1. Xi Y, Xu P. Global colorectal cancer burden in 2020 and projections to 2040. Transl Oncol. 2021;14(10):101174.

2. Dalisay DS, Tenebro CP, Sabido EM, Suarez AFL, Paderog MJV, Reyes-Salarda R, et al. Marine-Derived Anticancer Agents Targeting Apoptotic Pathways: Exploring the Depths for Novel Cancer Therapies. Mar Drugs. 2024;22(3).

3. Peng J, Yuan JP, Wu CF, Wang JH. Fucoxanthin, a marine carotenoid present in brown seaweeds and diatoms: metabolism and bioactivities relevant to human health. Mar Drugs. 2011;9(10):1806−28.

4. Nova P, Gomes AM, Costa-Pinto AR. It comes from the sea: macroalgae-derived bioactive compounds with anti-cancer potential. Crit Rev Biotechnol. 2024;44(3):462−76.

5. Tavares JO, Cotas J, Valado A, Pereira L. Algae Food Products as a Healthcare Solution. Mar Drugs. 2023;21(11).

6. Ji CF, Ji YB. Laminarin-induced apoptosis in human colon cancer LoVo cells. Oncol Lett. 2014;7(5):1728−32.

7. Zeng M, Wu X, Li F, She W, Zhou L, Pi B, et al. Laminaria Japonica Polysaccharides effectively inhibited the growth of nasopharyngeal carcinoma cells in vivo and in vitro study. Exp Toxicol Pathol. 2017;69(7):527−32.

8. van Weelden G, Bobinski M, Okla K, van Weelden WJ, Romano A, Pijnenborg JMA. Fucoidan Structure and Activity in Relation to Anti-Cancer Mechanisms. Mar Drugs. 2019;17(1).

9. Han YS, Lee JH, Lee SH. Antitumor Effects of Fucoidan on Human Colon Cancer Cells via Activation of Akt Signaling. Biomol Ther (Seoul). 2015;23(3):225−32.

10. Kim EJ, Park SY, Lee JY, Park JH. Fucoidan present in brown algae induces apoptosis of human colon cancer cells. BMC Gastroenterol. 2010;10:96.

11. Zhang Z, Teruya K, Eto H, Shirahata S. Fucoidan extract induces apoptosis in MCF-7 cells via a mechanism involving the ROS-dependent JNK activation and mitochondria-mediated pathways. PLoS One. 2011;6(11):e27441.

12. Newton K, Wickliffe KE, Dugger DL, Maltzman A, Roose-Girma M, Dohse M, et al. Cleavage of RIPK1 by caspase-8 is crucial for limiting apoptosis and necroptosis. Nature. 2019;574(7778):428−31.

13. Lin Y, Devin A, Rodriguez Y, Liu ZG. Cleavage of the death domain kinase RIP by caspase-8 prompts TNF-induced apoptosis. Genes Dev. 1999;13(19):2514−26.

14. Murakami Y, Miller JW, Vavvas DG. RIP Kinase-Mediated Necrosis as an Alternative Mechanism of Photoreceptor Death. Oncotarget. 2011;2(6):497−509.

15. Martens S, Bridelance J, Roelandt R, Vandenabeele P, Takahashi N. MLKL in cancer: more than a necroptosis regulator. Cell Death Differ. 2021;28(6):1757−72.

16. Xu X, Lai Y, Hua ZC. Apoptosis and apoptotic body: disease message and therapeutic target potentials. Biosci Rep. 2019;39(1).

17. Goldar S, Khaniani MS, Derakhshan SM, Baradaran B. Molecular mechanisms of apoptosis and roles in cancer development and treatment. Asian Pac J Cancer Prev. 2015;16(6):2129−44.

18. Di Cristofano F, George A, Tajiknia V, Ghandali M, Wu L, Zhang Y, et al. Therapeutic targeting of TRAIL death receptors. Biochem Soc Trans. 2023;51(1):57−70.

19. Pistritto G, Trisciuoglio D, Ceci C, Garufi A, D’Orazi G. Apoptosis as anticancer mechanism: function and dysfunction of its modulators and targeted therapeutic strategies. Aging (Albany NY). 2016;8(4):603−19.

20. Singh R, Letai A, Sarosiek K. Regulation of apoptosis in health and disease: the balancing act of BCL-2 family proteins. Nat Rev Mol Cell Biol. 2019;20(3):175−93.

21. Voss AK, Strasser A. The essentials of developmental apoptosis. F1000Res. 2020;9.

22. Morris G, Walker AJ, Berk M, Maes M, Puri BK. Cell Death Pathways: a Novel Therapeutic Approach for Neuroscientists. Molecular Neurobiology. 2017;55(7):5767−86.

23. Chan KWK, Chung HY, Ho WS. Anti-Tumor Activity of Atractylenolide I in Human Colon Adenocarcinoma In Vitro. Molecules. 2020;25(1).

